# Inhaled corticosteroids downregulate the SARS-CoV-2 receptor ACE2 in COPD through suppression of type I interferon

**DOI:** 10.1101/2020.06.13.149039

**Authors:** Lydia J Finney, Nicholas Glanville, Hugo Farne, Julia Aniscenko, Peter Fenwick, Samuel V Kemp, Maria-Belen Trujillo-Torralbo, Maria Adelaide Calderazzo, Jadwiga A Wedzicha, Patrick Mallia, Nathan W Bartlett, Sebastian L Johnston, Aran Singanayagam

## Abstract

Coronavirus disease 2019 (COVID-19) caused by SARS-CoV-2 is a new rapidly spreading infectious disease. Early reports of hospitalised COVID-19 cases have shown relatively low frequency of chronic lung diseases such as chronic obstructive pulmonary disease (COPD) but increased risk of adverse outcome. The mechanisms of altered susceptibility to viral acquisition and/or severe disease in at-risk groups are poorly understood. Inhaled corticosteroids (ICS) are widely used in the treatment of COPD but the extent to which these therapies protect or expose patients with a COPD to risk of increased COVID-19 severity is unknown. Here, using a combination of human and animal *in vitro* and *in vivo* disease models, we show that ICS administration attenuates pulmonary expression of the SARS-CoV-2 viral entry receptor angiotensin-converting enzyme (ACE)-2. This effect was mechanistically driven by suppression of type I interferon as exogenous interferon-β reversed ACE2 downregulation by ICS. Mice deficient in the type I interferon-α/β receptor (*Ifnar1*^−/−^) also had reduced expression of ACE2. Collectively, these data suggest that use of ICS therapies in COPD reduces expression of the SARS-CoV-2 entry receptor ACE2 and this effect may thus contribute to altered susceptibility to COVID-19 in patients with COPD.

## Introduction

Coronavirus disease 2019 (COVID-19) caused by SARS-CoV-2 infection is a new rapidly spreading infection which can cause a spectrum of disease ranging from a mild self-limiting illness to severe respiratory failure requiring ventilatory support. Current guidance advocates that high-risk individuals including those with chronic lung diseases such as severe asthma and COPD should be shielded to reduce risk of acquisition of the virus (1). This guidance is based upon extensive prior knowledge that these conditions are exquisitely susceptible to being exacerbated by a range of respiratory virus infections (2). However, early evidence has indicated that the prevalence of asthma and COPD among hospitalised COVID-19 cases may be lower than the general population, in contrast to other chronic comorbidities such as hypertension and diabetes, raising speculation of a possible protective phenotype (3). Conversely, COPD (but not asthma) has been shown to be associated with greater risk of COVID-19 related mortality (4, 5), suggesting that these patients could theoretically be protected from acquisition of the virus but paradoxically at increased risk of complications if they become infected.

Inhaled corticosteroids (ICS) are mainstay therapies for airways diseases and confer beneficial effects including protection against exacerbations (6, 7), suggesting that these drugs may reduce the risk of virus acquisition or alternatively suppress virus-induced inflammation and prevent symptomatic manifestations. Conversely, we and others, have shown that ICS have the adverse effect of suppressing innate immune responses to rhinovirus and influenza infection, leading to increased virus replication (8–10), although the opposite (protective) effect of ICS has been reported *in vitro* for the seasonal coronavirus 229E (11) and SARS-CoV-2 (12). It is thus unclear whether, overall, ICS impart a protective or detrimental effect on immune responses to SARS-CoV-2 and the extent to which these widely used drugs protect or expose patients with asthma or COPD to COVID-19 is unknown.

SARS-CoV-2 utilises the entry receptor Angiotensin-converting enzyme (ACE)-2 with priming of the serine protease TMPRSS2 (Transmembrane protease, serine 2) to gain entry into the respiratory mucosa and cause active infection (13). Increased epithelial ACE2 expression has been recently reported in smokers and subjects with COPD (14) and is postulated to be a factor predisposing these individuals to adverse outcome from COVID-19. Conversely, ACE2 is downregulated in asthma (15), an effect that may be due to suppressive effects of type 2 cytokines (16) or related to ICS use (17). Emerging evidence also indicates that ACE2 expression co-localizes with immune genes involved in interferon signalling pathways.(18) Moreover, Ziegler *et al* recently elucidated that *ACE2* is an interferon-stimulated gene in human respiratory epithelial cells (19), indicating that antiviral pathways may be important in regulation of pulmonary ACE2 expression. We have previously reported that ICS potently suppress epithelial expression of type I IFNs and interferon stimulated genes (ISGs) in a range of *in vitro* and *in vivo* COPD models (8) and it is plausible that ICS-mediated suppression of IFN might drive downregulation of ACE2 in the lungs and thus be an important determinant of susceptibility to SARS-CoV-2 in chronic lung disease patients.

Here, we show that ICS administration attenuates pulmonary expression of ACE2, an effect observed consistently across a range of human and animal COPD models. Using functional experiments, we demonstrate that the downregulation of ACE2 is mechanistically related to suppression of type I interferon by ICS. These data indicate that use of ICS therapies alter expression of the SARS-CoV-2 entry receptor and may thus contribute to altered susceptibility to COVID-19 in patients with COPD.

## Results

### *ACE2* mRNA expression is reduced in COPD patients taking inhaled corticosteroids

Recent data indicate that sputum expression of *ACE2* mRNA is reduced in asthmatic subjects taking ICS (17) but it is unclear whether similar suppression occurs in the context of COPD. We therefore initially used a community-based cohort of 40 COPD subjects (20) to determine whether ICS use affects ACE2 expression in COPD. 36 out of 40 subjects had sufficient sample for evaluation and were stratified according to current use (n=18) or non-use (n=18) of ICS. There were no significant differences between these groups in terms of age, disease severity, smoking status or other comorbidities known to affect ACE2 expression and/or associated with increased risk of COVID-19 (**Table 1**). Sputum cell *ACE2* mRNA expression was detectable in 22/36 COPD subjects (61.1%) and, consistent with prior observations in asthma (17), significantly reduced in ICS users compared to non-users (**Fig 1a**). Sputum cell expression of the serine protease *TMPRSS2* that is used by SARS-CoV-2 for mucosal entry (13) was detectable in all subjects with no significant difference observed between ICS users and non-users (**Fig 1b**). Similarly, the alternative SARS-CoV receptor CD147 (21) (gene: *Basigin BSG)* was detectable in all subjects with no difference observed between ICS users and non-users (**Supplementary Fig 1**).

**Table 1:** Demographic and clinical characteristics of COPD patients included in mRNA analyses in Figure 1 and Supplementary Fig 1

|  | ICS users<br>n=18 | ICS Non-users<br>n=18 | P value |
| --- | --- | --- | --- |
| **Age** | 68.5 (61.25 – 73.5) | 67 (62.25-70.75) | 0.61 |
| **Male** | 12 (66.7%) | 14 (77.8%) | 0.71 |
| **FEV1 (% predicted)** | 65 (50-77) | 67 (63-82) | 0.43 |
| **GOLD stage I/II/III/IV** | 3/11/2/2 | 5/10/3/0 | - |
| **Prior Exacerbations in preceding year** | 1.5 (1-4) | 1 (0-1) |  |
| **Current smoker** | 6 (33.3%) | 7 (38.9%) | 1.0 |
| **Comorbidities** |  |  |  |
| Diabetes | 1 (5.6%) | 2 (11.1%) | 1.0 |
| Hypertension | 2 (11.1%) | 0 (0%) | 0.49 |
| Obesity | 3 (16.7%) | 2 (11.1%) | 1.0 |
| Ischaemic heart disease | 3 (16.7%) | 2 (11.1%) | 1.0 |
| **Inhaled corticosteroid type** | Fluticasone 8 (44.4%)<br>Budesonide 9 (50%)<br>Beclomethasone 1 (5.6%) | n/a | - |

**Figure 1:**
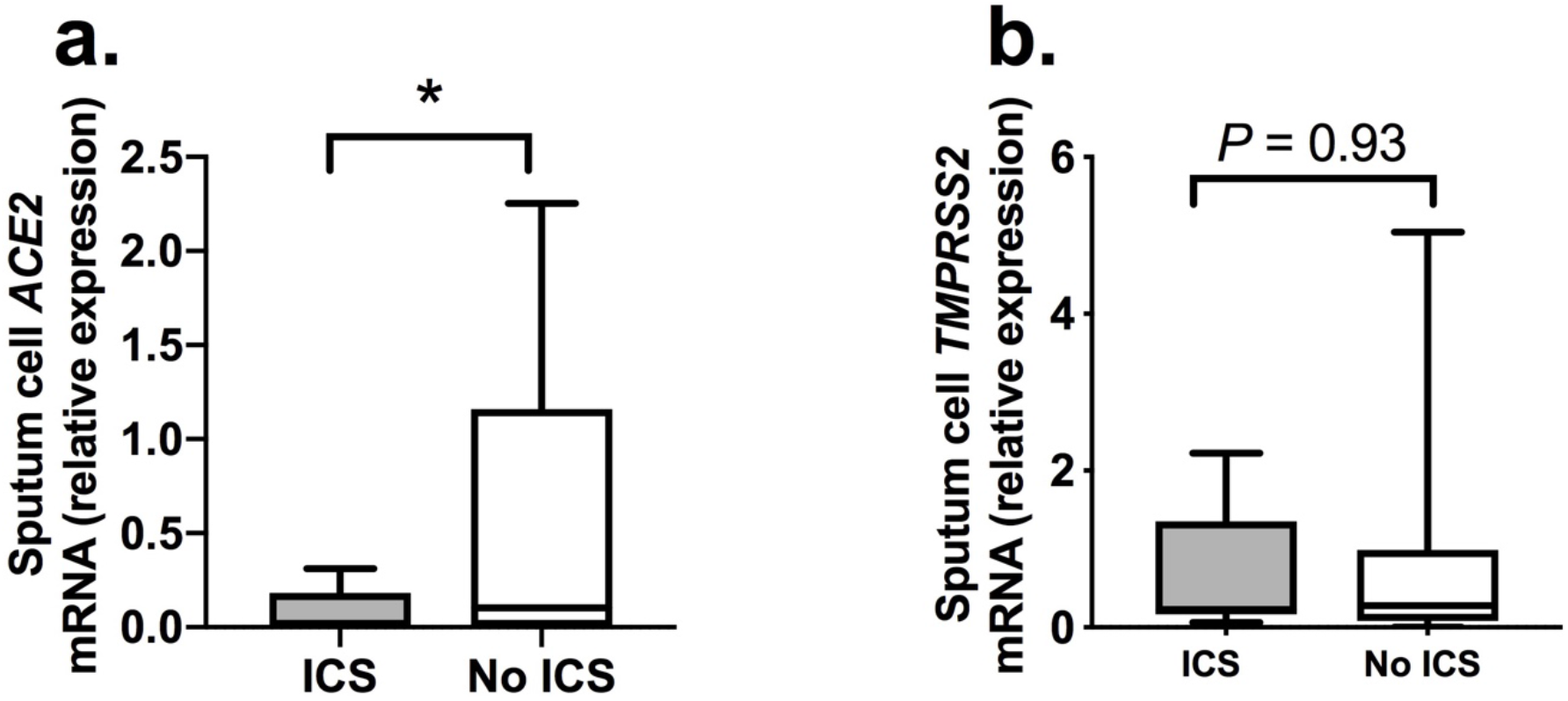
Sputum gene expression of *ACE2* and *TMPRSS2* in COPD subjects stratified according to use or non-use of inhaled corticosteroids. Sputum samples were taken from a cohort of patients with COPD when clinically stable for at least 6 weeks. Patients were stratified according to current use or non-use of inhaled corticosteroids (ICS). Sputum cell mRNA expression of (a) *ACE2* and (b) *TMPRSS2* was measured by quantitative PCR. Box-and-whisker plots show median (line within box)), interquartile range (IQR; box) and minimum to maximum (whiskers). Statistical comparisons made using Mann Whitney *U* test. \**P*<0.05

### Inhaled corticosteroid administration suppresses pulmonary expression of ACE2 in mice

Given that cause and effect cannot be inferred from a cross-sectional human study, we next evaluated whether experimental pulmonary administration of the ICS fluticasone propionate (FP) in mice, at a dose previously shown to induce lung glucocorticoid receptor (GR) activation (8, 22), had similar effects on *Ace2* expression. A single administration of 20μg FP in mice (**Fig 2a**) induced significant downregulation of pulmonary *Ace2* mRNA expression at 8 hours (a timepoint where we have also previously shown that significant GR activation occurs (8)). This effect persisted at 24 hours post-administration but had resolved from 48 hours onwards (**Fig 2b**). Consistent with effects observed in human sputum, FP administration had no effect expression of *Tmprss2* or *Bsg* in mouse lung (**Supplementary Fig 2**). Suppression of *Ace2* by FP occurred in a dose-dependent manner with loss of suppression at a ten-fold lower concentration (2μg)(**Fig 2c**), a dose at which effects on GR activation are also lost (8). We observed similar suppression of lung *Ace2* mRNA expression with administration of 20μg of other commonly used ICS budesonide and beclomethasone, suggesting that the effect of ICS on *Ace2* is not class dependent (**Fig 2d**). To corroborate the effects observed on *Ace2* mRNA expression, we subsequently measured protein levels in lung homogenate of ICS-treated mice by ELISA. We observed similar suppression of total lung ACE2 protein occurring at 24 hours post-administration, consistent with effects observed at the mRNA level (**Fig 2e**).

**Figure 2:**
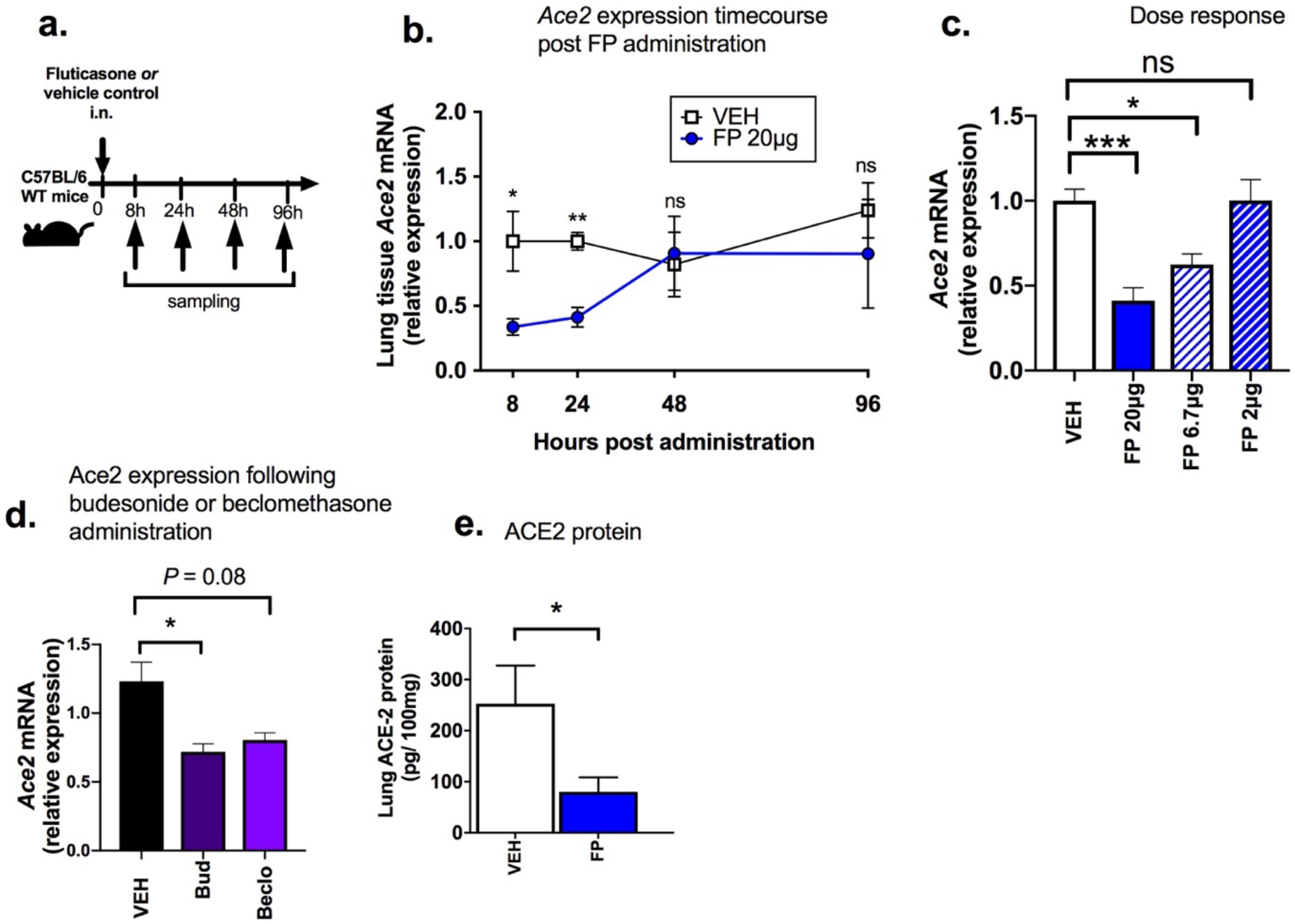
Inhaled corticosteroid administration downregulates ACE2 expression in mouse lung. (a) C57BL/6 mice were treated intranasally with a single dose of fluticasone propionate (FP) or vehicle DMSO control. (b) Lung *Ace2* mRNA expression was measured by qPCR at the indicated timepoints following 20μg FP administration (c) Lung *Ace2* mRNA expression was measured by qPCR at 24 hours following single dose administration of FP at 20, 6.7 and 2μg doses. (d) Lung *Ace2* mRNA expression was measured by qPCR at 24 hours following single dose administration of 20μg budesonide (Bud), beclomethasone (Beclo) or vehicle control. (e) ACE2 protein in lung tissue homogenate measured by ELISA at 24 hours following single dose administration of 20μg FP. Data represents mean (+/−) SEM of 4-5 mice per treatment group, representative of at least two independent experiments. Data analyzed by one way ANOVA with Bonferroni post-test. \**P*<0.05 *** *P* <0.001. ns = non-significant. VEH=vehicle control.

### Downregulation of ACE2 by FP is functionally related to suppression of type I IFN

ACE2 has recently been reported to co-localise with expression of type I-IFN related genes (13, 18) and has also been shown to be an interferon stimulated gene (ISG) in the respiratory tract (19), suggesting that type I IFN may be a major regulator of pulmonary ACE2 expression. Consistent with this, we observed that basal lung *Ace2* expression in mice positively correlated with mRNA expression of the ISG 2’-5’ *OAS* mRNA and BAL concentrations of IFN-λ in a combined analysis of FP and vehicle treated mice (**Fig 3a**). Given our previous data showing that COPD patients treated with ICS have reduced basal airway expression of *IFNβ (8)*, we hypothesized that downregulation of ACE2 by FP may be functionally related to its suppressive effects on type I IFN signalling. Accordingly, recombinant IFN-β administration (**Fig 3b**) could reverse FP-mediated suppression of *Ace2* mRNA and ACE2 protein (**Fig 3c**), indicating that the effect of FP on ACE2 expression is functionally related to suppressive effects on type I IFN.

**Figure 3:**
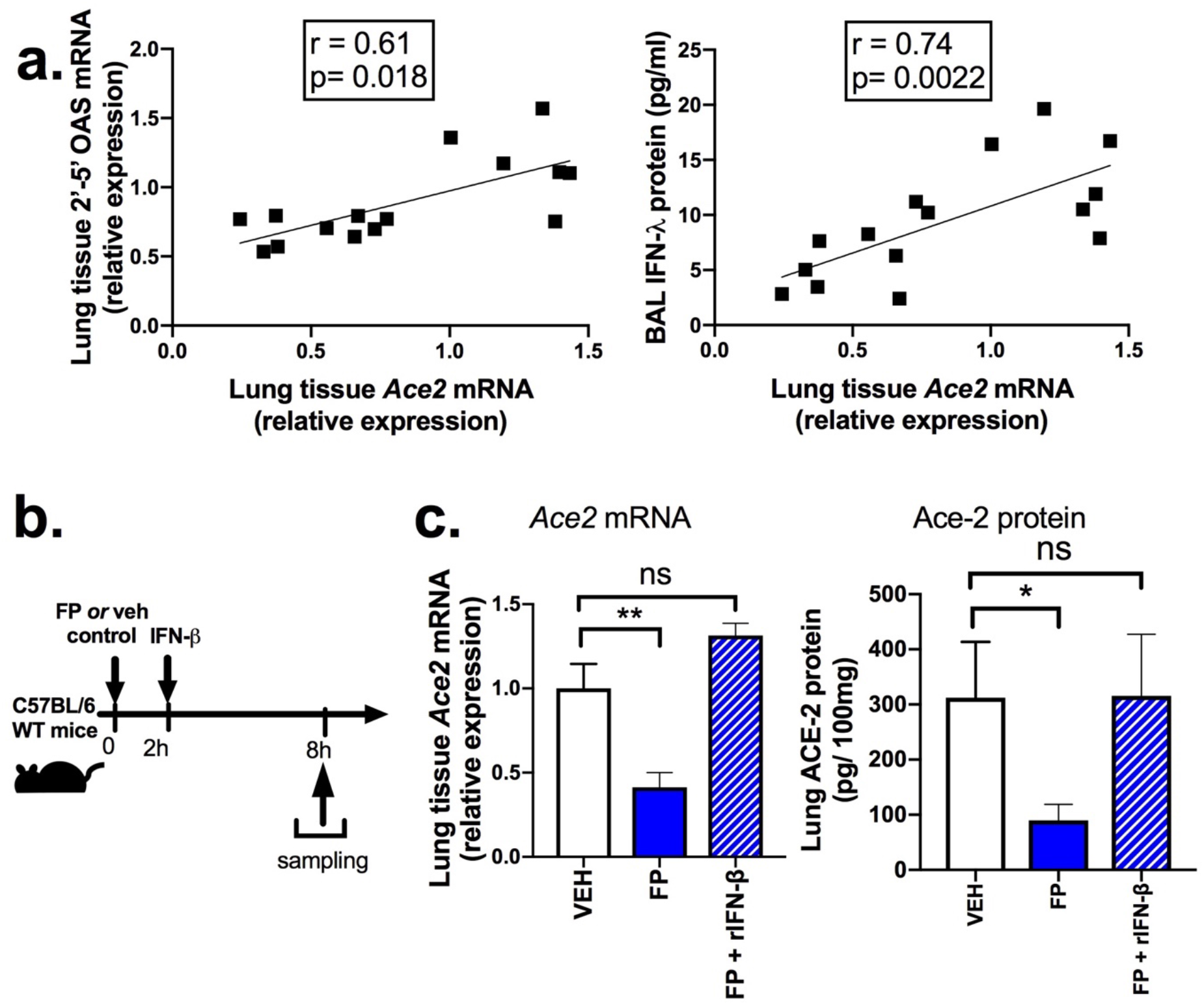
Downregulation of ACE2 by inhaled corticosteroid is functionally related to suppression of type I IFN. (a) Correlation between lung *Ace2* mRNA and lung *2’-5’OAS* and BAL IFN-λ in C57BL/6 mice. (b) C57BL/6 mice were treated intranasally with fluticasone propionate (20μg) or vehicle DMSO control and additionally with recombinant IFN-β or PBS control. (c) Lung *Ace2* mRNA expression and (d) Lung ACE2 protein concentrations were measured at 8 hours post FP administration. Data represents mean (+/−) SEM of 5 mice per treatment group, representative of at least two independent expreriments. Data analysed by Spearman’s rank correlation (a) or one way ANOVA with Bonferroni post-test (c). \**P*<0.05 ** *P* <0.01. ns = non-significant.

### *Ifnar ^−/−^* mice have reduced basal expression of ACE2

To further confirm the functional importance of type I IFN in regulating pulmonary ACE2, we evaluated basal pulmonary expression levels in mice deficient in IFN signalling (*Ifnar*^−/−^). Compared to wild-type control mice, *Ifnar*^−/−^ mice had a small, but statistically significant, reduction in lung *ACE2* mRNA expression (**Fig 4a**) with a concomitant trend (*P*=0.15) towards reduced lung ACE2 protein levels (**Fig 4b**). These observations further confirm the key regulatory role played by type I IFN signalling in pulmonary expression of ACE2.

**Figure 4:**
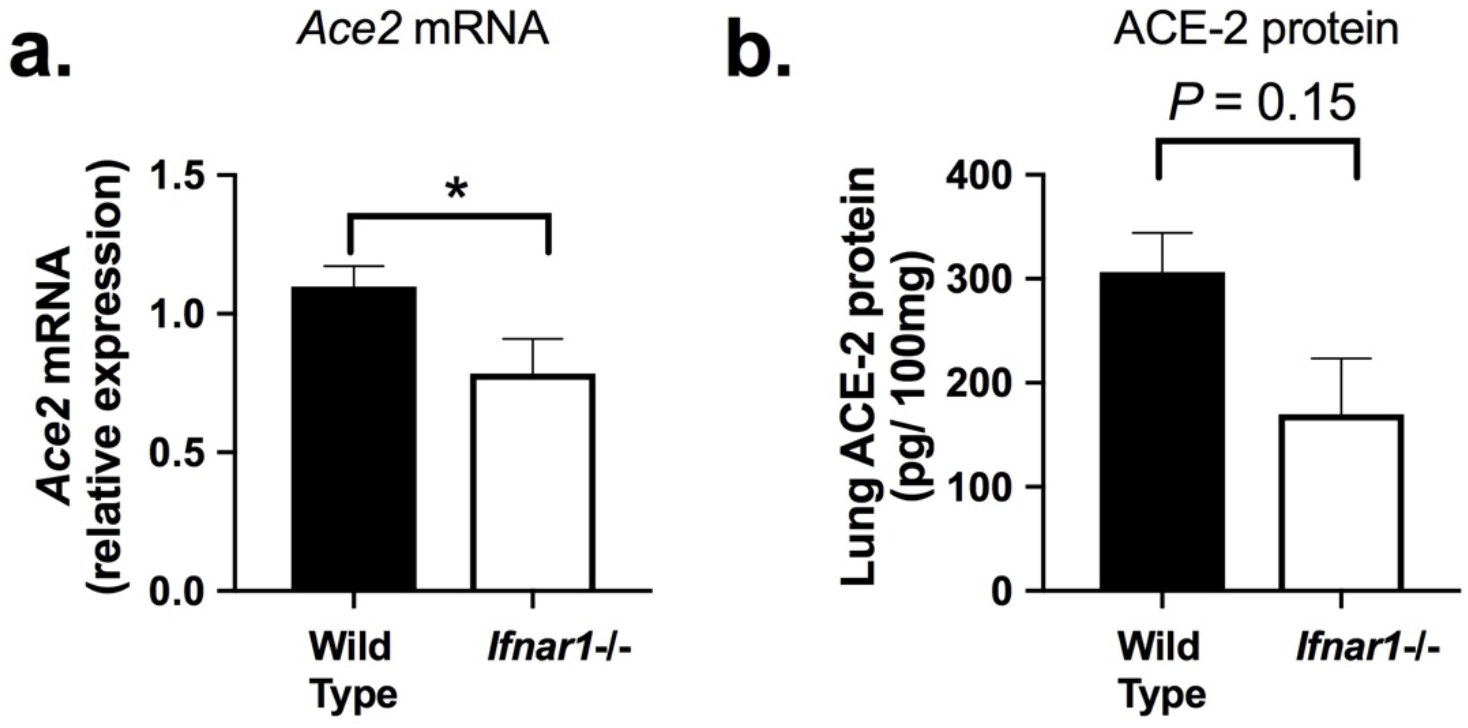
Type I IFN receptor deficient mice have reduced pulmonary ACE2 expression. Lung tissue was harvested from wild type or *Ifnar1*−/−C57BL/6 mice. (a) Lung *Ace2* mRNA expression was measured by qPCR and (b) Lung ACE2 protein concentration was measured by ELISA. Data represents mean (+/−) SEM of 4-5 mice per treatment group, representative of at least two independent expreriments. Data analysed by T test. * *P* <0.05 ** *P* <0.01.

### The suppressive effect of ICS upon *ACE2* expression occurs at the bronchial epithelium

Existing data indicate that ACE2 is expressed primarily in the nasal and bronchial epithelium and is absent from immune cells (16). Given our prior data indicating that FP also exerts its inhibitory effects on immunity principally at the pulmonary epithelium (8, 23), we next assessed whether suppressive effects on ACE2 were also observed following *ex vivo* ICS administration in cultured COPD bronchial epithelial cells (BECs, **Fig 5a**). Baseline characteristics of the subjects included in these analyses are shown in table 2. In keeping with recent *in situ* expression studies in COPD patients (14), we found that basal expression of *ACE2* was increased by ~3 fold in BECs from COPD patients compared to healthy non-smokers (**Fig 5b**). Consistent with our findings in human ICS users and in the mouse model of ICS administration (**Figs 1&2**), FP administration (at a clinically relevant concentration of 10nM) induced ~75% suppression in *ACE2* expression (**Fig 5c**).

**Table 2:**
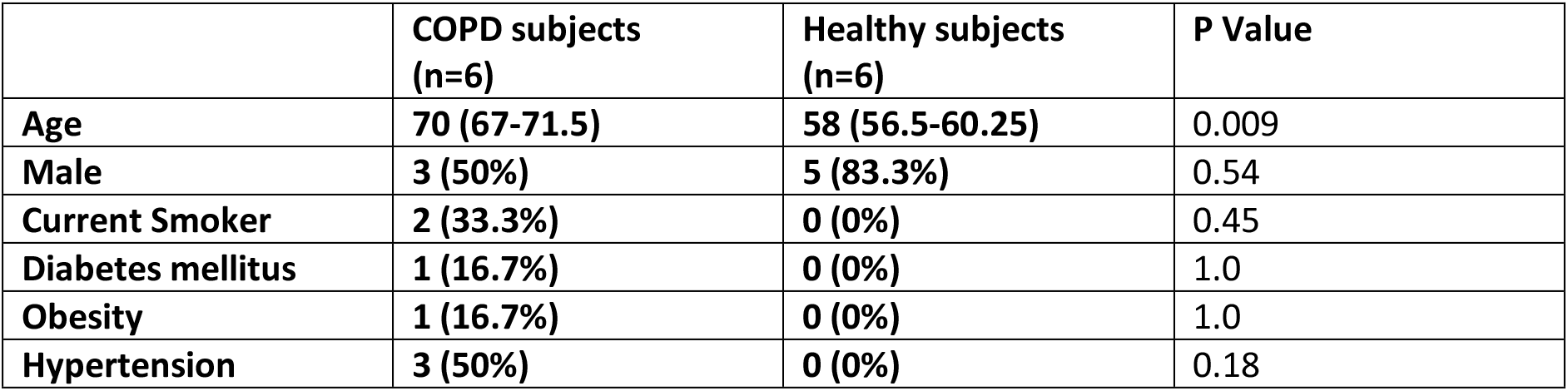
Demographic and clinical variables in COPD and healthy subjects included in primary airway epithelial cell experiments (Figure 5)

**Figure 5:**
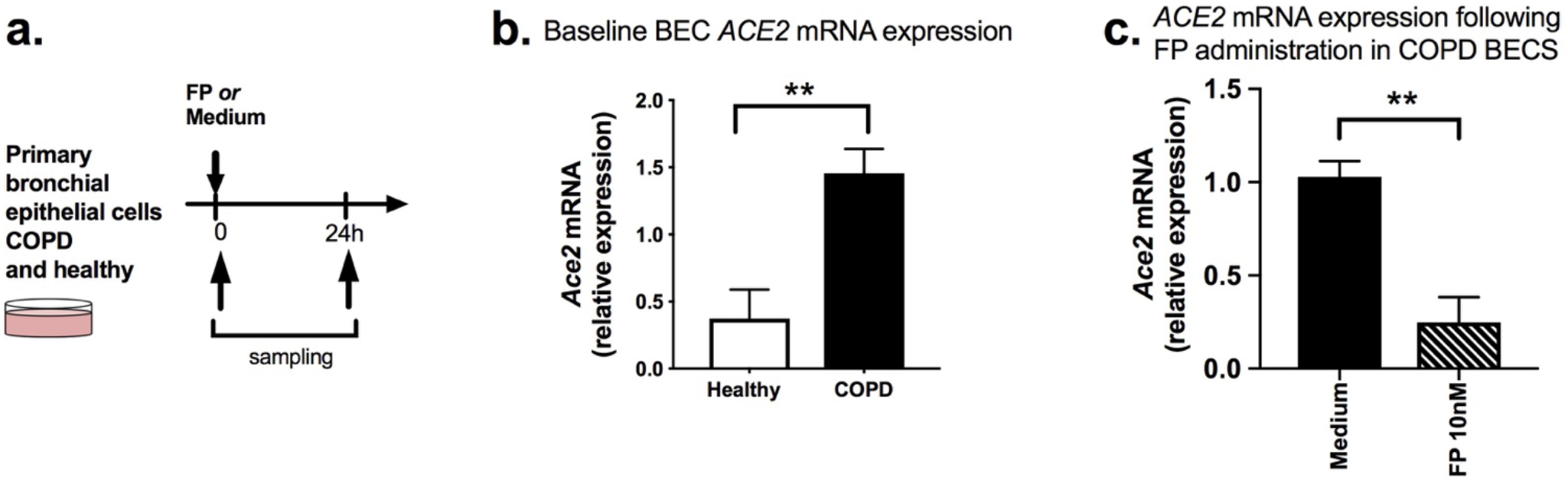
ACE2 expression is increased in COPD and suppressed by fluticasone administration in cultured bronchial epithelial cells. (a) Primary bronchial epithelial cells from 6 COPD subjects and 6 healthy control subjects were cultured *ex vivo* and treated with 10nM fluticasone propionate (FP) or medium control. Cell lysates were collected. ACE2 mRNA expression was measured at baseline (b) and at 24h after FP administration (c) in COPD BECs. Data shown as median (+/−IQR). Statistical comparisons made using Mann Whitney *U* test. \*\**P*<0.01.

### FP downregulates pulmonary ACE2 expression in a mouse model of COPD

To further confirm that ICS administration suppresses ACE2 expression in COPD, we employed a mouse model of elastase-induced emphysema which recapitulates many hallmark features of human disease (24). Mice were treated intranasally with a single dose of porcine pancreatic elastase (**Fig 6a**). and lung ACE2 expression was measured at 10 days after administration (timepoint at which COPD-like disease features are established (24)) and a further 7 days later. In keeping with our findings in human COPD cells, elastase-treated mice had significantly increased (~5-fold) lung *Ace2* expression at 10 days with further enhancement to >15-fold at 17 days (**Fig 6b**). Administration of a single dose of FP at 10 days attenuated the significant upregulation of lung *Ace2* mRNA (**Fig 6c**) and ACE2 protein concentrations measured 24 hours later in elastase-treated mice (**Fig 6d**). Therefore, the suppressive effects of ICS on ACE2 also occur in an *in vivo* model of COPD-like disease.

**Figure 6:**
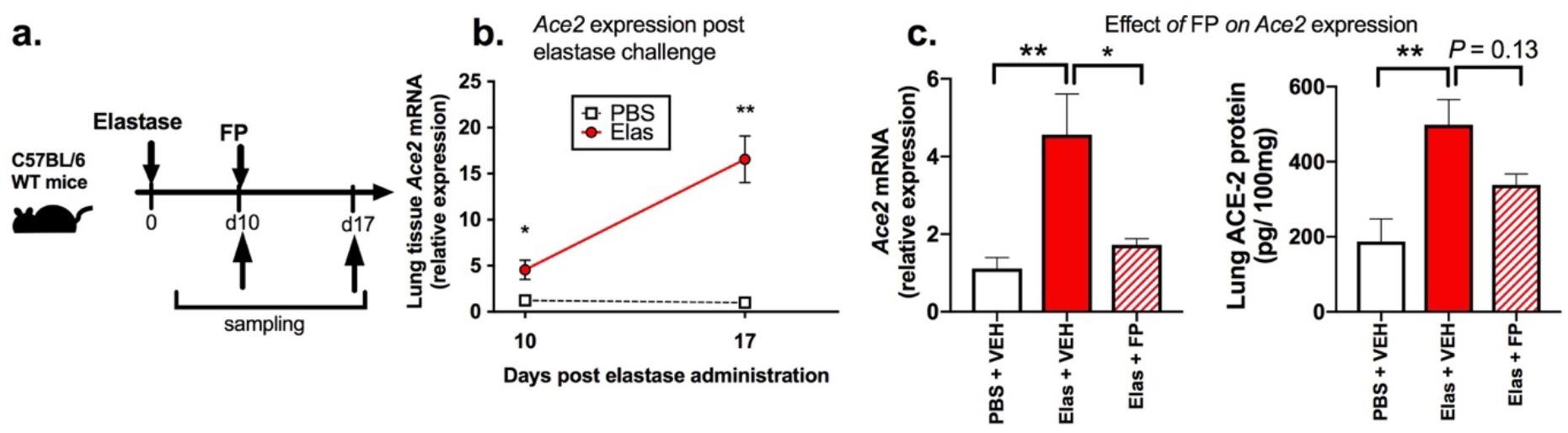
ACE2 expression is increased in a mouse model of COPD and suppressed by fluticasone administration. (a) C57BL/6 were treated intranasally with a single dose of elastase or PBS as control and lung tissue harvested at 10 or 17 days later. Some mice were treated with fluticasone propionate (FP) of vehicle (VEH) control at 10 days, before sampling 24 hours later. (b) Lung *Ace2* mRNA expression in elastase- versus PBS-treated mice at 10 and 17 days after treatment. (c) At 10 days after elastase treatment, C57BL/6 mice were treated with a single dose of 20μg fluticasone propionate (FP) or vehicle (VEH) control. Lung tissue was harvested at 24 hours after FP administration. *ACE2* mRNA was measured by qPCR (left panel) and ACE2 protein was measured by ELISA (right panel). Data represents mean (;/-) SEM of 5 mice per treatment group, representative of at least two independent experiments. Data analysed by one way ANOVA with Bonferroni post-test. \**P*<0.05 \*\**P*<0.01. ns = non-significant.

## Discussion

The mechanisms driving altered susceptibility to COVID-19 in chronic lung diseases and whether the commonly used therapies ICS promote or protect against infection by SARS-CoV-2 is a crucial question for the field. In this study, we demonstrate consistently, across a range of human and animal models, that ACE2, a receptor that facilitates entry of SARS-CoV-2 in the respiratory tract, is upregulated in COPD and suppressed by ICS treatment. Our studies indicate a novel mechanism for downregulation of ACE2 by ICS through suppression of type I interferon.

There is clear evidence to support the premise that ACE2 mediates cell entry of SARS-CoV2 into the respiratory tract and also acts as a major receptor for SARS-CoV1 and NL63 coronaviruses (13, 25, 26). Viral entry occurs as a two-step process with initial binding of the N-terminal portion of the viral protein to the ACE2 receptor, followed by viral protein cleavage facilitated by the receptor transmembrane protease serine 2 (TMPRSS2)(13). Therapeutic blockade of TMPRSS2 inhibits entry of SARS-CoV-1 and −2 into cells, supporting a critical role for this protease in viral pathogenesis (13). Our data indicates clear downregulation of ACE2 by ICS but with no concomitant effect on TMPRSS2. Although the precise role of ACE2 in the pathogenesis of COVID-19 is not fully characterised, several lines of evidence indicate the important role it plays in mediating virus entry and facilitating disease pathogenesis of SARS-CoVs. In SARS-CoV-1, greater ACE2 expression increases *in vitro* susceptibility to infection (27). Moreover, in animal models, overexpression of *Ace2* enhances SARS-CoV-1 entry, anti-ACE2 antibodies can block viral invasion and reduced pulmonary pathology is observed in *Ace2-* deficient mice (25, 28, 29).

ACE2 is expressed primarily in the nasal goblet cells and type II pneumocytes within the respiratory tract (30) and is upregulated in subject groups known to be associated with increased disease severity including elderly individuals (31) and patients with diabetes (17), indicating that it may play a clinically important role in governing susceptibility to virus acquisition and/or development of severe disease in at-risk groups. ACE2 expression has similarly been shown to be increased in COPD: using combined transcriptomic and immunohistochemical analyses, Leung *et al* recently demonstrated that epithelial *ACE2* expression is increased in bronchial brushings/tissue samples from COPD subjects versus healthy controls (32), effects also previously shown in cigarette smoke exposure animal models (33). Our data showing increased *ACE2* expression in cultured airway epithelial cells and in an elastase mouse model of COPD are consistent with these findings. Rates of patients with COPD being hospitalized with COVID-19 have been relatively low, ranging from 1.1% in Chinese cohorts to 5% in US cohorts (34–37) However, patients with COPD have greater risk of severe disease and mortality from COVID-19 (4). These data suggest that COPD may be associated with possible protection against the need for hospitalisation (possibly due to reduced risk of virus acquisition) but increased propensity to severe disease upon infection. The original SARS-CoV-1 pandemic was also characterised by an extremely low prevalence of chronic lung disease comorbidities (38, 39), an effect that could also have been driven by ICS-mediated suppression of ACE2 in these subjects. The mechanism underlying these putative alterations in susceptibility has not been extensively explored. Our data suggests that suppression of ACE2 by the commonly used therapies ICS may be one important factor that dictates susceptibility in COPD.

Currently, a direct causal link between ACE2 and increased susceptibility to acquisition of SARS-CoV-2 or subsequent severity has not been proven and we cannot conclude unequivocally that downregulation of ACE2 by ICS is an effect that would confer protection clinically. In asthma, a disease where ICS are more commonly prescribed than in COPD, *ACE2* expression is reduced compared to healthy subjects (15) and also further attenuated in ICS users (17). In contrast to COPD, asthma has not been shown to be associated with increased COVID-19 mortality (5) and the more widespread use of ICS with associated suppression of ACE2 could theoretically be one factor driving this. Conversely, It is important to note that there is evidence to suggest that downregulation of ACE2 could also theoretically worsen outcome. In mouse models of experimentally-induced acid aspiration and sepsis, genetic deletion of *Ace2* worsens acute lung injury, an effect that is partially rescued by recombinant ACE2 administration(40). ACE2 also degrades angiotensin II which can drive production of pro-inflammatory cytokines (41, 42) which may be detrimental in the context of the hyper-inflammation that is characteristic in severe COVID-19 (43). Downregulation of ACE2 by ICS could therefore remove critical homeostatic protective functions within the lungs and thereby promote severe disease in COVID-19.

ICS use can also impart a number of other detrimental effects on innate immunity to other respiratory viruses (which may also occur in the context of SARS-CoV-2 infection) including suppression of type-I interferon leading to increased virus replication (8-10) and virus-induced pathology including mucus hypersecretion and secondary bacterial infection (8). We therefore cannot currently ascertain whether the clear suppressive effect of ICS upon ACE2 expression shown in our experiments would overall have protective or detrimental effects in the context of clinical disease. Further functional manipulation experiments (e.g. SARS-CoV-2 challenge in ICS treated ACE2 depleted animals) coupled with prospective studies of ICS use as a predictor of susceptibility to SARS-CoV-2 infection are required to understand the direct implications of this effect. If ACE2 downregulation does confer protection against SARS-CoV-2 infection then our results would suggest that these therapies should be continued stringently in subjects with asthma and COPD. It may also indicate that therapies that are currently in trials, such as recombinant IFN-β, could theoretically induce the adverse effect of increasing ACE2 and promoting greater viral entry.

Understanding the mechanism through which ICS suppress expression of ACE2 is important to delineate how this effect could either be harnessed as a protective factor or reversed if deemed to be detrimental. We have previously shown that ICS can suppress type I IFN both at steady state and during active virus infection (8). Here, we show that this effect directly drives suppression of ACE2 expression since administration of recombinant IFN-β in combination with FP could reverse the downregulation of ACE2. Furthermore, *Ifnar* ^−/−^ mice which lack type I IFN signalling also had reduced ACE2 providing additional evidence that expression is directly regulated by type I IFN. These findings are consistent with recent studies showing that genes relevant to IFN pathways (*IFNAR1*, *IFITM1*) are expressed in *ACE2*+ type 2 pneumocytes and that type I IFNs can upregulate ACE2 in a range of experimental systems (13). We additionally showed that two other commonly used ICS, budesonide and beclomethasone, have similar effects on ACE2, confirming that, in contrast to impairment of anti-bacterial immunity (22), the suppressive effect of ICS on ACE2 is not class-dependent. This is also consistent with the mechanism of suppressed IFN as we have also shown previously that budesonide suppresses IFN to a similar degree to FP (8).

In summary, these studies indicate that inhaled corticosteroid use in COPD suppresses the expression of the SARS-CoV-2 receptor ACE2 through a type I interferon-dependent mechanism. These effects are likely to contribute to altered susceptibility to COVID-19 in patients prescribed these therapies. Further studies are now needed to elucidate the precise effects that altered ACE2 expression has on the acquisition of SARS-CoV-2 infection and associated severity in COVID-19.

## Materials and Methods

### The St. Mary’s Hospital COPD cohort

A cohort of 40 individuals previously recruited for a longitudinal study carried out at St Mary’s hospital between June 2011 and December 2013 was used to investigate the relationship between ICS use and ACE2 expression. The study received ethical approval from the East London Research Ethics Committee (study number 11/LL/0229). All included subjects had spirometrically confirmed COPD and were seen when clinical stable (no episodes of acute infections, antibiotic treatment or oral corticosteroid treatment within the previous 8 weeks). All patients underwent clinical assessment, spirometry and had spontaneous or induced sputum, taken and processed as previously described.(20, 44) Subjects were stratified on the basis of current use or non-use of ICS, either in a single agent inhaler or in combination with bronchodilators.

### Treatment of cultured primary bronchial epithelial cells

Primary BECs were obtained bronchoscopically from six subjects with spirometrically confirmed COPD and six healthy non-smoking control subjects. The study was approved by the Bromley ethics committee (REC: 15/LO/1241). Primary cells were cultured in collagen coated T75 flasks in LH-9 medium until 80% confluent before being seeded at 2.5 × 10^5^ cells per well in a 24-well plate. Cells were treated with 10nM FP (Sigma-Aldrich) or vehicle dimethyl sulfoxide (DMSO), as previously reported (23).

### Mouse models

8-10 week old female C57BL/6 mice were used for all studies. Mice were purchased from Charles River laboratories UK, housed in individually ventilated cages within specific pathogen free conditions. *Ifnar*^−/−^ mice were bred in house on a C57BL/6 background. All animal experiments were carried out under the authority of the UK Home Office within the Animals (Scientific Procedures) Act 1986 (project licence number 70/7234).

Fluticasone propionate, Budesonide or Beclomethasone powder (Sigma-Aldrich) was resuspended in dimethyl sulfoxide (DMSO) at a concentration of 357 μg/mL and then diluted 1: 1000 in sterile PBS. Mice were treated intranasally under light isofluorane anaesthesia with a 50μL solution containing 20, 6.7 or 2μg of fluticasone or 20μg of budesonide or beclomethasone (8, 22). Mice were culled for endpoint analyses at 8, 24, 48 or 96 hours after administration of FP. In some experiments, two hours after FP administration, mice were additionally treated intranasally with 50μL of PBS containing 10^4^ units of recombinant IFN-β (8).

### RNA extraction, cDNA synthesis and quantitative PCR

RNA was extracted from cell lysates of primary airway epithelial cells, human sputum cells or mouse lung tissue using an RNeasy kit (Qiagen). 2μg was utilised for cDNA synthesis using the Omniscript RT kit (Qiagen). Quantitative PCR was carried out using previously described specific primers and probes and normalized to 18S rRNA housekeeping gene. (8) Reactions were analysed using the ABI 7500 real time PCR machine (Applied Biosystems).

### Protein assays

A commercially available ELISA duoset kit (Abcam) was used to measure total ACE2 concentrations in mouse lung homogenate. The lower limit of detection for this assay is 10 pg/mL. IFN-λ concentrations in mouse BAL were also measured using a commercially available ELISA duoset kit (Bio-techne), as previously reported (8).

### Statistical analyses

For animal experiments, group sizes of 5 mice per condition were used and data is presented as mean+/−SEM representative of at least two independent experiments. Data were analysed by one-way ANOVA with Bonferroni’s multiple comparison test. mRNA expression in sputum cells of ICS users versus non users or in cell lysates of *ex vivo* cultured epithelial cells treated with FP was compared by Mann Whitney *U* test. All analyses were performed using GraphPad Prism version 8. Differences were considered statistically significant when *P*<0.05.

## AUTHOR CONTRIBUTION

**Conception and Design:** LJF, NG, JAW, NWB, PM, SLJ, AS; **Experimental Work:** LJF, NG, HF, JA, PF, SVK, MT, MAC, AS; **Writing and Critical Evaluation of Manuscript:** All authors

## ACKNOWLEDGEMENTS

Nill

---

**Supplementary Figure 1:**
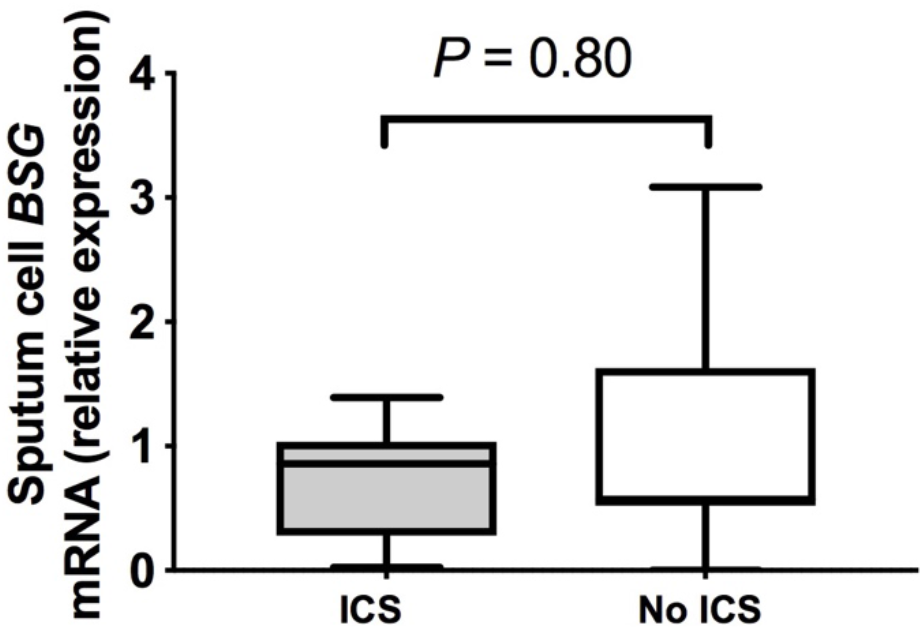
Sputum gene expression of *BSG* in COPD subjects stratified according to inhaled corticosteroid use. Sputum samples were taken from a cohort of patients with COPD when clinically stable for at least 6 weeks. Patients were stratified according to current use or non-use of inhaled corticosteroids (ICS). Sputum cell mRNA expression of *BSG* was measured by quantitative PCR. Box-and-whisker plots show median (line within box)), interquartile range (IQR; box) and minimum to maximum (whiskers). Statistical comparisons made using Mann Whitney *U* test.

**Supplementary Figure 2:**
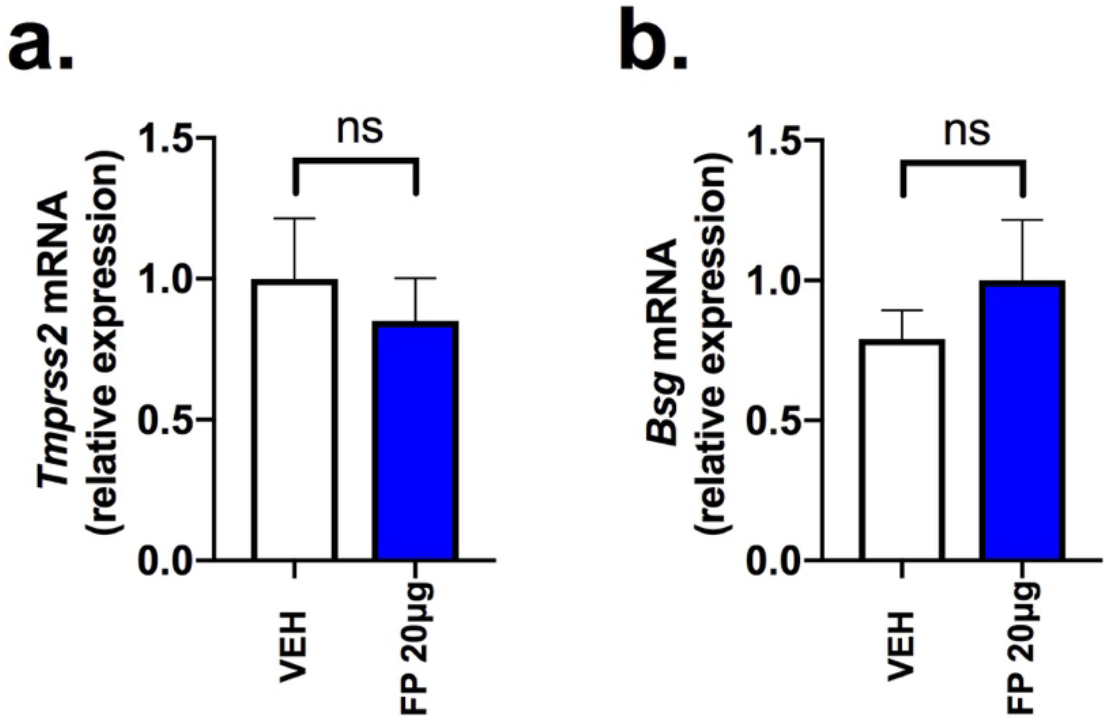
No effect of fluticasone propionate administration on expression of *Tmprss2* or *Bsg* in mouse lung. C57BL/6 mice were treated intranasally with a single 20μg dose of fluticasone propionate (FP) or vehicle DMSO control. (a) lung *Tmprss2* and (b) lung *Bsg* mRNA expression was measured by qPCR at 8 hours following FP administration. Data represents mean (+/−) SEM of five mice per treatment group, representative of at least two independent expreriments. Data analysed by one way ANOVA with Bonferroni post-test. \**P*<0.05 \*\*\**P*<0.001. ns = non-significant.

